# A method for creating custom 3D-printed molds to facilitate zebrafish imaging studies, including of cardiac development

**DOI:** 10.1101/2025.06.01.657246

**Authors:** James Christian Miller, Prashanna Koirala, Maria Fernanda Argote de la Torre, Marjan Farsi, Jaret Lieberth, Rabina Shrestha, Joshua Bloomekatz

**Author notes:** Corresponding author –.

## Abstract

Embryo mounting is one of the technical challenges researchers encounter when undertaking an imaging project. Embryos need to be oriented in a reproducible manner such that the tissue of interest is accessible to a microscope objective for the entire imaging period. To overcome this challenge researchers can embed embryos in viscous media or create specialized dishes and casts to hold embryos in a desired orientation during imaging. Here we describe a method for using a cheap stereolithographic (SLA) 3D-printer to manufacture re-usable molds that create agarose wells in which embryos can be mounted for imaging. These agarose wells provide a reliable means for orienting multiple embryos for imaging. This method includes a design framework that can be easily customized for a variety of tissues, organisms and imaging challenges. Using this method we have created molds for imaging cardiac development in zebrafish for both upright and inverted microscopes. By utilizing materials and equipment that are accessible this method allows researchers to easily create molds specific to their mounting needs.

**SUMMARY:** Here we describe a method that uses a cheap stereolithographic (SLA) 3D printer to create molds to facilitate the reproducible mounting of zebrafish embryos for imaging studies. Using this method, we’ve created molds for imaging cardiac morphogenesis in zebrafish embryos.

## INTRODUCTION

Visualizing cellular dynamics within an intact animal is essential for comprehending the pathologies of congenital diseases and the fundamental processes of development and homeostasis. This visualization encompasses imaging at multiple spatial scales to identify physiological dynamics across an entire organism as well as at the tissue, cellular and sub-cellular levels. And it involves imaging modalities ranging from immunofluorescence studies of fixed specimens to time-lapse imaging of live specimens. These imaging modalities require holding intact specimens in specific orientations, such that a region of interest is accessible to the microscope objective. Zebrafish embryos are an ideal model organism for imaging studies due to their optical transparency, genetic tractability, external fertilization and vertebrate origin.

However, mounting zebrafish embryos in a specific orientation in a consistent and reproducible manner has represented a continual challenge. Mounting is especially challenging for the visualization of heart development in zebrafish embryos because cardiac morphogenesis occurs in different planes^1^.

Multiple approaches have been developed to address the challenges of mounting embryos for imaging. One of the simplest methods involves mounting zebrafish embryos between two coverslips with layers of tape in between the coverslips to prevent crushing. This method allows for imaging on both inverted and upright microscopes^2^. Zebrafish embryos can also be mounted in a highly viscous media or gel such as methylcellulose or 0.1-0.8% low melting point agarose in order to hold embryos in specific orientations^2–4^. These low-throughput approaches work well for many imaging projects, however ensuring embryo integrity and reproducibly obtaining specific orientations can be troublesome. To increase imaging throughput and facilitate mounting, regularly spaced wells can be created from solidified agarose in an imaging dish^5–8^. To facilitate the creation of these agarose wells, plastic molds have been created^9–11^. While these molds can be applied to imaging along the standard dorsal-ventral-lateral axes^10,12^, they are often made for specific imaging challenges, such as for use in 96 well plates^13^ or for the detection of the first heartbeat^14^. In specialized cases entire imaging dishes have been created^5,15,16^. For example, a microfluidic chamber (named ZEBRA) made-out of PDMS facilitates both imaging and exposure of embryos to chemicals^17^. These molds are often made from silicone or machined from lucite^10,18,19^. However the advent of relatively cheap three-dimensional (3D) printers has democratized the creation of customized molds that are used to create agarose imaging wells.

3D printing, also called additive or layered manufacturing, involves producing a 3D object from a digital design by the sequential addition and fusion of individual layers^20,21^. Stereolithography (SLA) utilizes ultraviolet (UV) light to cure a light sensitive liquid resin in sequential layers^22^. SLA printers are cost-effective, fast, highly accurate and can be used with a variety of resins including those that deliver a smooth surface^16^. The process begins with computer-aided design (CAD) software, where digital models of the object can be designed de-novo or modified from pre-existing designs. The digital model is then virtually sliced into printable layers. Using these digital layers a 3D printer constructs the object, layer by layer. Here we present a method and framework for the custom design and creation of imaging molds using a cheap commercially available SLA 3D-printer. Additionally, we use this framework to create molds for imaging different stages of zebrafish cardiac development on either inverted or upright microscopes and for imaging zebrafish embryos along their lateral, dorsal or ventral axes.

### PROTOCOL

To create agarose wells of different shapes in which embryos can be oriented for imaging, we developed a design that utilizes a base attached to pegs (see Fig. 1A). The shape of these pegs can be customized to orient embryos of different developmental stages and to focus on different tissues. Below we describe a protocol for customizing these pegs and for using this system to create agarose wells.

**Figure 1:**
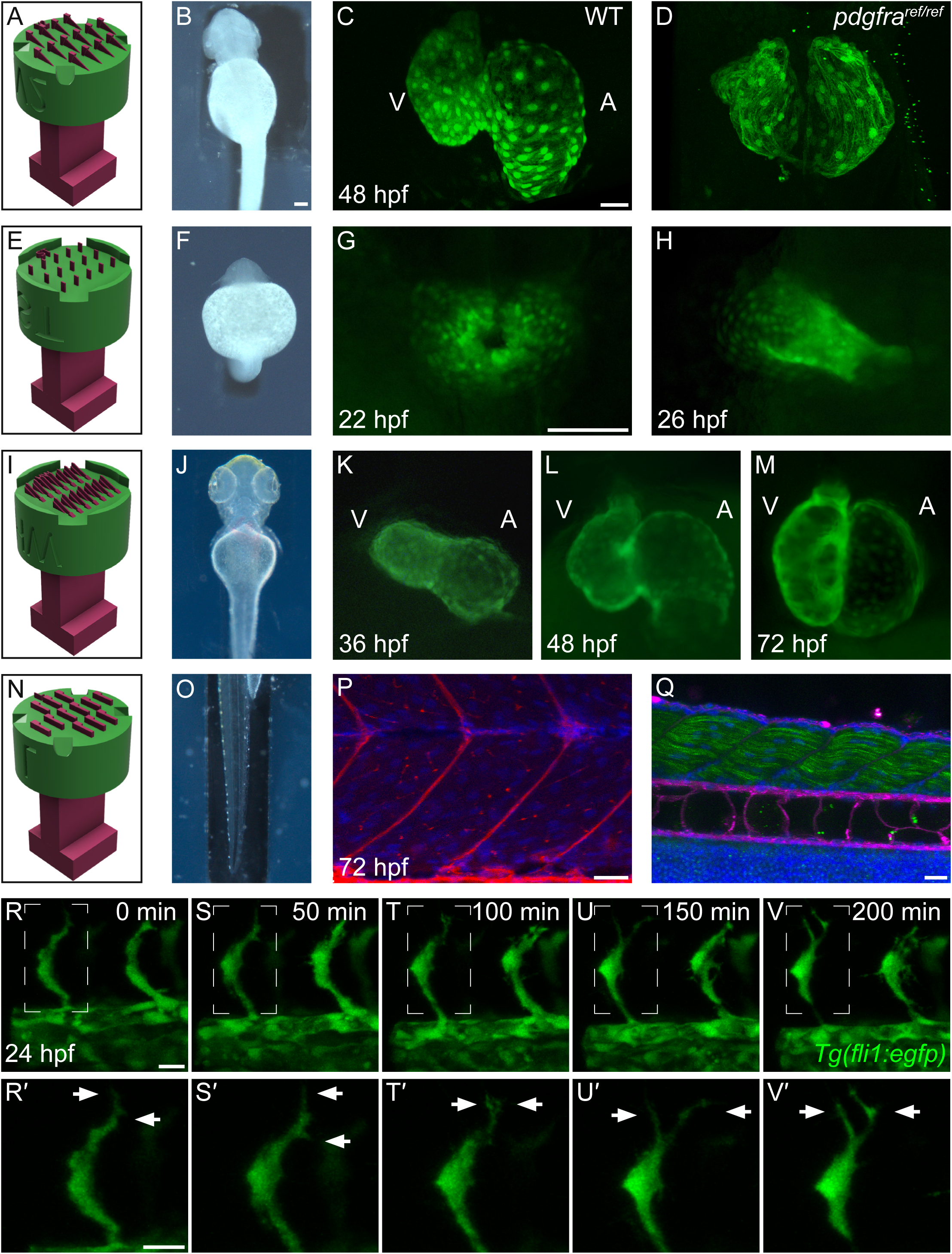
3D-printed molds can be customized for specific tissues and imaging challenges. A-V’. We have designed and printed molds to facilitate the mounting and imaging of different zebrafish tissues including the heart (A-M) and the trunk (N-V’). These molds are useful for multiple different imaging experiments including analyzing mutant embryos with an inverted confocal microscope (A-D), comparing cardiac morphology at different developmental timepoints with an upright stereo microscope (E-M), immunofluorescence analysis of the sub-cellular localization of glycans (N-Q), and live time-lapse studies (R-V’). **A, E, I, N** Design schematics of 3D-printed molds. **B, F, J, O** shows zebrafish embryos mounted in agarose wells created by the 3D-printed molds shown in A, E, I, N, respectively. **C, D** Three-dimensional reconstructions of *Tg(myl7:egfp)* transgenic wild-type and *pdgfra^ref/ref^*mutant hearts at 48 hours post-fertilization (hpf) created with an inverted confocal microscope. **G, H, K-M** Developmental time-series of early zebrafish heart development, captured with an upright stereo microscope, using the *Tg(myl7:egfp)* transgene (green). Two different molds ST (G, H) and HW (K-M) were used due to the changing position of the zebrafish heart. **P, Q** Three-dimensional reconstructions of a zebrafish trunk stained with rhodamine-labeled lectins – ricinus communis agglutinin (RCA-120, P) and wheat germ agglutinin (WGA, Q), which preferentially bind to terminal β-linked galactose containing glycans or GlcNAc and sialic acid containing glycans, respectively (red in P, magenta in Q). Embryos were co-stained with DAPI (P, Q – blue) and the F59 antibody revealing fast fiber skeletal muscle (Q – green). **R-V** Timepoints from a time-lapse movie (see Supplemental movie 1) of intersomitic vessel (ISV) formation created using the *Tg(fli1a:egfp)* transgene. **R’-V**’ are magnifications of the boxed regions in R-V, showing filopodia formation. The L2 mold was used to create agarose wells for P,Q and R-V. RCA-120 and WGA from Vector labs (catalog #RLK-2200, 1:300 dilution), F59 from DHSB (1:50), Goat anti-mouse Alexa-488 from Thermofisher (catalog #A28175, 1:300). V, ventricle, A, atrium. Scale bars: C, D = 30μm; G, H, K, L, M = 100 μm; P = 20 μm; Q = 30 μm; R-V’= 15 μm.

#### A. Creating a custom-designed imaging mold

1. **Sketch:** Create a preliminary sketch of the specimen or embryo positioned in the desired orientation. It is essential to consider the desired embryo orientation and whether the microscope you are going to use contains upright or inverted objectives. Additional considerations include the size of the dish, the number of embryos to be mounted and the spacing between them.
2. **Measure**: Measure the dimensions of the sample (eg. length, width and height) using a microscope combined with a software tool such as FIJI^23^.
3. **Design pegs:** Identify a shape that corresponds to the specimen’s geometry. Then using the measurements from step 2 in combination with a computer-aided design (CAD) software such as TinkerCAD (Autodesk Inc., US.) or Autodesk Fusion 360 (Autodesk, Inc., US.) create a peg that approximates the shape and size of the sample. This shape can be created denovo using TinkerCAD which provides a variety of pre-designed shapes which can be combined and overlapped or using Autodesk Fusion which allows for full customization of the shape. Additionally, one can modify one of our pre-designed pegs (Supplemental file A1). Tip: Tapering the peg as in Fig. 1A, can help to orient embryos to ensure the tissue of interest is closest to the objective.
4. **Combine peg design with base design:** In TinkerCAD or Autodesk Fusion add your custom designed pegs to one of our pre-designed bases (Supplemental files A2-3). The pegs are then be duplicated and spaced evenly to allow for high-throughput imaging. Ensure all pegs reach the same height away from the base. We have created two bases: one for imaging with an upright microscope and one from imaging with an inverted microscope. The base for imaging with an upright microscope has an outer edge that prevents the pegs from sinking all the way to bottom (for example, Fig. 1E). However, for imaging with an inverted microscope the pegs need to touch the bottom so that the sample can be as close to the coverslip-bottom as possible (for example, Fig. 1N). Additional features of the bases include divots that allow agarose to spread evenly between the pegs and a label positioned on one side of the bottom of the base. The asymmetry of the label allows one to uniquely identify each peg position and thus each mounted embryo.
5. **Prepare the design for 3D printing**: Position critical components such as the pegs away from the printer stage to minimize the risk of warping. Transfer the customized mold design or one of the pre-designed molds (Supplemental files B1-13) to a slicer software such as Chitubox (CBD-Tech, China), which converts the design into commands for the 3D printer. Supports are also added at this stage to ensure that delicate aspects of the mold don’t break during the printing. Multiple pieces can be printed simultaneously by either using a single STL file containing all molds or arranging individual STL files together using a software such as Chitubox.
6. **Printing:** SLA printers are our preferred 3D printer-type for these imaging molds due to their feasibility, precision and their creation of smooth surfaces. We have used a Elegoo Mars Pro 2 printer with a water washable resin (Elegoo P05BB028). After printing, the molds are washed in water, supports are removed and then the molds are placed under UV light to ensure the solidification of the resin. This can be performed using the Elegoo Mercury V3.0 or by washing in the sink and placing the molds under UV light for example from a tissue-culture hood. Note: Gloves and a lab coat should be worn when handling uncured resin and cleaning the molds, since uncured resin can be a skin irritant. Note: SLA printers use UV light to cure the resin, which can be harmful to the eyes and skin. Ensure that the print chamber is securely enclosed and avoid looking directly at the UV light source.

#### B. A procedure for using 3D-printed molds to create agarose wells

1. Agarose wells are made with 2% agarose in E3 media (1.578 mM NaCl, 0.169 mM KCl, 0.249 mM CaCl_2_*2H_2_O, 0.161 mM MgSO_4_*7H_2_O). Place a mold in the center of a coverslip bottom dish with the arrow pointing down towards the well (see Fig 2A-B). Pipette 750 µl of 2% molten agarose/E3 along half of the imaging mold’s perimeter (see Fig 2C). Tip: Molten agarose should be non-viscous but cool enough to pipette without creating bubbles. Note: We use 60mm dishes with a 30mm coverslip bottom well. For molds designed for upright microscopes a standard 60mm dish is fine. Note: A slightly different protocol is used for the screening mold (see below).
2. Pipette another 750 µl of 2% molten agarose/E3 along the other half of the imaging mold’s perimeter (see Fig 2D).
3. Wait 5 to 7 minutes for the agarose to solidify, observing the transition from a clear liquid to an opaque blue gel (see Fig 2E).
4. Place the outer ring (Suppl. Fig. 2B) over the agarose mold assembly with the arrow pointed towards the agarose (see Fig 2F).
5. While gently holding down the outer ring mold, carefully pull straight up on the inner mold to detach it from the agarose (see Fig 2G). Tip: To avoid breaking or damaging the agarose wells, don’t push down too hard on the outer ring and don’t twist or shift the mold when pulling upwards.
6. Remove the outer ring, revealing the agarose wells in the coverslip bottom dish (Fig. 2H).
7. Inspect the agarose wells under a microscope to confirm correct sizing and production. Tip: For imaging on an inverted microscope, a thin film of agarose often needs to be lifted off the bottom of the well. This can be done with forceps.

### Modified protocol for using the screening mold

1. Pour molten 2% agarose/E3 into the petri dish until it is approximately half-way full.
2. Hold the mold with the pegs facing upwards and pipet molten 2% agarose/E3 onto the pegs, until they covered (∼ 3 ml). Tip: Ensure that bubbles are not introduced by releasing only 90% of the molten agarose that has been drawn into the pipet.
3. Using a tilting motion, place one end and then the whole mold with pegs face down into the petri dish containing molten agarose (Suppl. Fig 3).
4. Allow the agarose to solidify for 5-7 mins.
5. Carefully remove the mold by using the handle to pry up one end and then the other end.

#### C. Mounting embryos in agarose wells with and without low-melting temperature agarose

1. Preparation: Dechorionate and anesthetize embryos in a 200 μg/ml tricaine solution prior to mounting. An ∼400 μg/ml tricaine solution is required to stop the heart from beating. (The protocols, experiments and findings described here were conducted in accordance with animal welfare regulations and guidelines, and the Institutional Animal Care and Use Committee at the University of Mississippi, protocol #24-008).
2. **a.** Without using low melting temperature agarose: Fill the imaging dish with 1x E3 and tricaine (200 μg/ml), add dechorionated anesthetized embryos to the imaging dish, use forceps to place and orient embryos in the agarose wells and then image (see Fig. 2I, J). **b.** Using low melting temperature agarose: While the molds are designed to place embryos in specific orientations, using low melting temperature agarose can ensure the embryos don’t drift during imaging or while you are transporting the imaging dish with the mounted embryos to the microscope. The steps below are derived from our previous protocol^6^.

**i.** Create tubes of low melting agarose: Dissolve 0.1-0.8% low-melting agarose in 5 ml E3 media and heat to 65-70 °C in a water bath. Occasionally invert the tube until the low-melting temperature agarose has dissolved. Aliquot the low-melting temperature agarose solution into microfuge tubes and place on a heat block set to 42 °C. Allow the solution to equilibrate to 42 °C for at least 1 hour. (It is generally advisable to use a freshly made low-melting temperature agarose solution.)
**ii.** Using a wide-bore glass pasteur pipette, transfer a single embryo into a tube containing the low-melting agarose solution. To avoid adding excess liquid to the low-melting temperature agarose solution, only transfer the embryo and liquid immediately surrounding it. Immediately transfer the embryo, which is now in low-melting agarose, to one of the agarose wells. Return the low-melting temperature agarose solution to the heat block for re-use.
**iii.** Using forceps, position the embryo (as shown in diagrams) in the well.
**iv.** Repeat steps i-iii for the other wells. (If the previous wells begin to dry out, a p200 pipetman can be used to add a small amount of E3 (∼50-100μL) on top of the well.)
**v.** Once embryos have been mounted in all wells and the low-melting temperature agarose has solidified, using a pipetman slowly add the tricaine/E3 solution to the imaging dish.
3. After imaging, embryos can be carefully extracted from the wells using forceps and allowed to develop further to ensure imaging did not cause developmental defects. The agarose wells can be removed from the imaging dish, which can be rinsed and reused. Rinsing with commercially available glass cleaner (eg. Windex) and then with Milli-Q water helps to remove salt-deposits on the coverslips.

## REPRESENTATIVE RESULTS

We used the above protocol to design molds for imaging different stages of cardiac development in zebrafish, including cardiac fusion, heart tube formation, cardiac looping and chamber formation. Since cardiac development doesn’t occur along the major anatomical axes of the embryo, we have customized the shape of the pegs for different developmental stages in order to properly position the embryos. For example, the anterior inner placement of the zebrafish heart at 48 hours post-fertilization (hpf) relative to the spherical yolk requires a sloped embryo orientation which we facilitated by creating pegs with tapered edges in mold V2 (Fig. 1A, B). Utilizing mold V2 to create agarose wells for imaging with an inverted confocal microscope we used the *Tg(myl7:egfp)* transgene^24^ to compare the cardiac morphology between a wild-type embryo and a severe *pdgfra^ref/ref^* mutant embryo^25^ displaying cardia bifida at 48 hpf (Fig. 1C, D). To image early events in heart development such as cardiac fusion and lumen formation, using an upright stereo microscope we designed a mold (named Standing, ST) that creates a well for the zebrafish trunk (Fig. 1E-H). This places the dorsal anterior side of the embryo directly under the objective of an upright microscope, allowing for these early stages of heart development to be visualized. During later stages of heart development, the heart is positioned on the ventral side of the embryo. For imaging these stages with an upright microscope, we designed a mold (named – Halfway, HW) that creates a shallow trough that holds embryos on their dorsal side for imaging cardiac looping and chamber development (Fig. 1I-M). Altogether we have designed and printed three molds for imaging cardiac development with an inverted microscope (Fig. 1A, Suppl. Fig. 1B-D) and two molds for imaging cardiac development with an upright microscope (Fig. 1E, 1I, Suppl. Fig. 1E-F).

**Figure 2:**
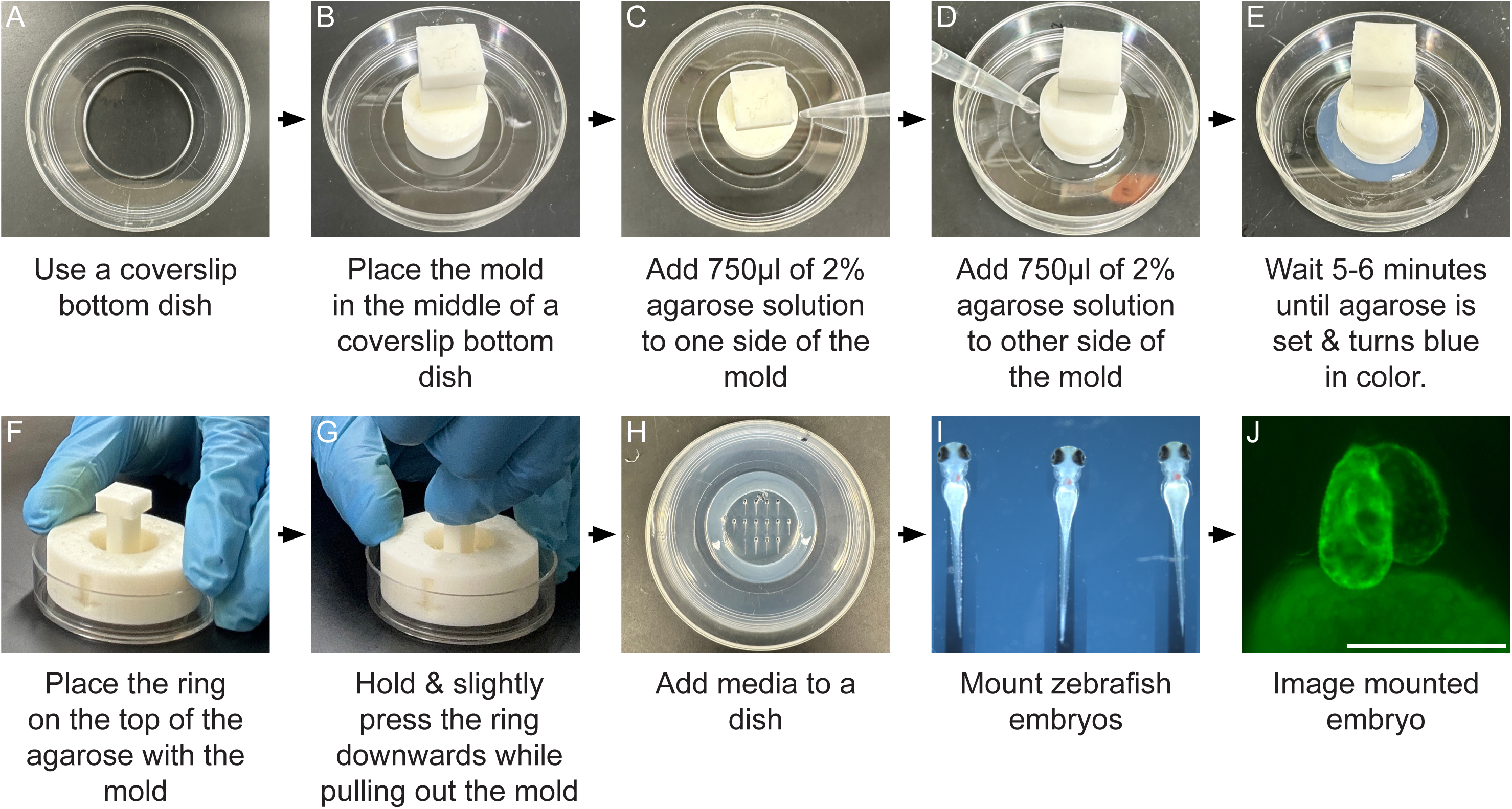
Protocol for using a 3D-printed mold to make agarose wells in a coverslip bottom dish. **A)** Use a 30mm coverslip bottom dish. **B)** Place the SLA 3D-printed mold in the center of the coverslip bottom dish. **C)** Slowly release 750μL of molten 2% agarose around the right half of the mold perimeter. **D)** Repeat Step C for the left half of the mold perimeter, evenly distributing a total of 1.5 ml of molten 2% agarose around the entire perimeter of the mold**. E)** Wait 5-6 minutes for the agarose to solidify (it turns slightly blue in color). **F)** Place the 3D-printed outer ring directly over the top of the inner mold and solidified agarose with the arrow pointing down. **G)** Remove the inner mold from the solidified agarose by gently pushing down on the outer ring while using the handle to simultaneously gently pull directly up on the inner mold. Be careful not to damage the agarose wells by twisting or shifting the mold as you pull up. **H)** Add enough media to the dish to cover the agarose wells. **I)** Mount and properly orient embryos in the agarose wells using a pipette and forceps. **J)** A representative image of a 96 hpf mounted embryo containing the *Tg(myl7:egfp)* transgene is shown. Scale in (J) = 100 μm.

Additionally, we created molds for imaging zebrafish embryos along the lateral, dorsal or ventral axes of the embryo with an inverted microscope. This includes a mold that orients zebrafish embryos in a lateral position for visualizing the zebrafish trunk (Fig. 1N, O). With this mold we imaged zebrafish trunks stained with the fluorescently conjugated lectins Ricinus Communis Agglutinin (RCA-120) and Wheat Germ Agglutinin (WGA) which preferentially bind to terminal β-linked galactose and N-acetylglucosamine (GlcNAc) or sialic acid containing glycans^26^, respectively. We observed that galactose containing glycans as detected by RCA-120 staining are enriched in myoseptal junctions (Fig. 1P), whereas either GlcNAc or sialic acid containing glycans as detected by WGA staining are enriched in the notochord (Fig. 1Q). WGA can bind to both GlcNAC and sialic acid containing glycans^26–29^. We also used the lateral mold to conduct time-lapse studies of intersomitic blood vessel (ISV) formation (Fig. 1R-V’).

Using the *Tg(fli1a:eGFP)* transgene^30^ we observed filopodia formation (Fig. 1R-V’, Supplemental movie 1), as previously reported^31^. These general molds (see Suppl. Fig. 1A, G – I) are designed for the imaging of 10-96 hpf zebrafish embryos along the ventral, dorsal and lateral axes.

We have also designed a mold for routine phenotypic and transgenic screening of 24-72 hpf embryos with an upright microscope (Suppl. Fig. 2A). And we have designed a coverslip holder for the mounting technique that utilizes two coverslips^2^ (Suppl. Fig. 2C). We created the coverslip holder with a fused deposition modeling (FDM) printer because these printers create stiffer lightweight products and the smooth surfaces created by SLA printers was not needed for this holder^32^.

## DISCUSSION

In this protocol we outline a method for designing 3D-printed molds that create agarose wells which can facilitate the mounting of embryos or tissues for imaging studies. The agarose wells made by these custom molds can be used for a range of imaging applications including immunofluorescence studies of fixed embryos and live timelapse imaging. And these molds can be made for imaging with either upright or inverted microscopes.

This method utilizes a foundation that contains two pieces: an inner plunger-shaped mold with pegs at the bottom and an outer ring-shaped mold. We have incorporated several key design features into these molds. The most significant of these features are the pegs located at the base of the inner mold. The shape of these pegs can be customized to create differently shaped agarose wells, which help to orient embryos in a reproducible manner for imaging. Additional design features include an impression of the mold name at one end of the agarose wells, creating an asymmetry that allows each well to be uniquely identified. And divots were added around the base of the inner mold to facilitate the innervation of molten agarose around the pegs.

Furthermore, while having pegs bottom-out at the coverslip is advantageous for inverted microscopes since it allows embryos to be placed as close to the coverslip as possible, this is not helpful for upright microscopes. Thus, for upright microscopes we added a lip around the bottom of edge of the inner mold. Finally, an outer ring placed over the inner mold after the agarose has solidified ensures that the agarose wells stay stuck to the imaging dish and that they aren’t damaged when the inner mold is removed.

Building on this foundation, we designed and tested molds for imaging cardiac development and for general zebrafish imaging. The molds for imaging cardiac development include 4 molds designed to be used with an inverted microscope - All (A), Cardiac fusion (CF), Lumen formation (LF) and Ventral 2 (V2); and two molds to be used with an upright microscope – Standing (ST) and Halfway (HW). The All mold creates a well for imaging cardiac specification from 0 – 10 hpf (Suppl. Fig. 1A). The CF mold helps embryos balance up-side down so that the process of cardiac fusion can be imaged from the dorsal side (Suppl. Fig. 1B). A steep gradient on one wall of the agarose well is created with the LF mold allowing lumen formation, which occurs under the head and on the side of the embryo, to be imaged (Suppl. Fig. 1C). For the V2 mold, a slope is created by the mold so that the head is tilted forward over the yolk, placing the heart which is on the ventral side of the embryo starting at 36 hpf perpendicular to the coverslip (Suppl. Fig. 1D). For imaging with an upright microscope such as a stereo microscope, the ST mold creates slender agarose wells in which to place the trunk. This stably positions the dorsal side of the embryo up towards an upright objective allowing heart development to be visualized from 14-28 hpf (Suppl. Fig. 1E). To image cardiac development on an upright microscope when the heart is on the ventral side of the embryo, we designed a mold that creates shallow agarose wells with a gradual incline (Suppl. Fig. 1F). We also designed molds for mounting zebrafish embryos for inverted microscopes along the major axes: dorsal (Suppl. Fig 1G), lateral (Suppl. Fig. 1H) and ventral (Suppl. Fig. 1I). For routine transgenic and mutant screening of zebrafish embryos with a fluorescent stereo microscope, we designed a large mold that creates 120 shallow agarose wells in which 24-72 hpf embryos can fit (Suppl. Fig. 2A). This mold is designed for an 100mm diameter dish. For traditional mounting using two coverslips, we designed a cheap coverslip holder. This coverslip holder fits most microscope stages and helps in the movement of embryos to and from the microscope (Suppl. Fig. 2C).

As we have optimized the procedure using these molds to create agarose wells, we have identified several key steps and tips for their successful design and use (See Table 1). For example, after the molds have been manufactured it is important to cure them with UV light, this can be done with a commercially available device (eg. Elegoo Mercury v3.0) or by leaving the molds under tissue culture hood with the UV light turned on. Additionally, it is important to verify that the 2% agarose solution is completely molten when adding it to the mold on the coverslip-bottom dish and that the right amount is added. If too much molten agarose is added the mold becomes difficult to extract from the solidified agarose. However, holes can appear in the agarose when too little molten agarose is added. We also found that when imaging on an inverted microscope it was often necessary to use forceps to peel a thin layer of agarose off the bottom of the well, this allows the embryo to get as close to the coverslip as possible.

**Table 1:**
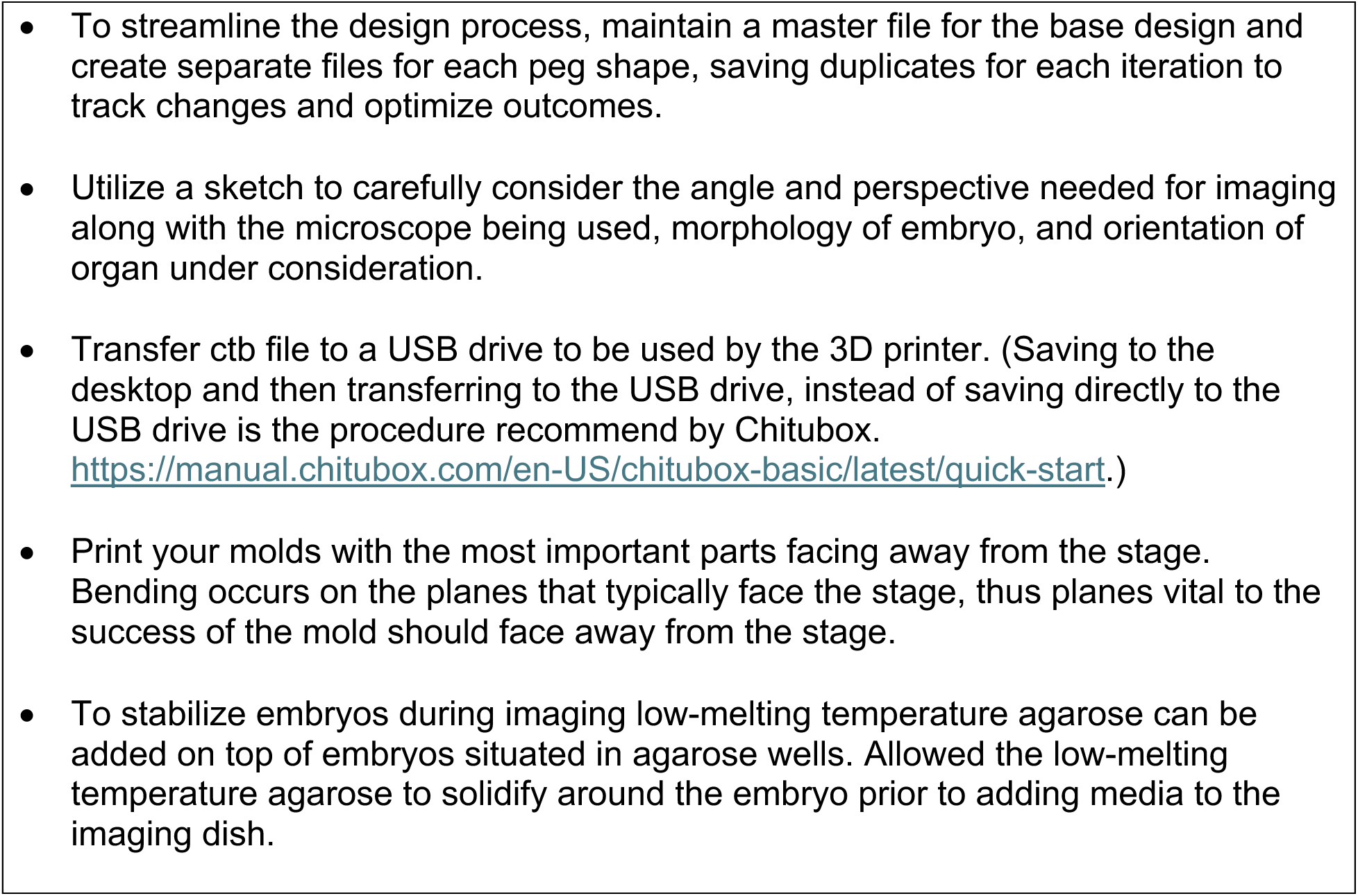
Tips for designing and creating custom molds for mounting zebrafish embryos.

Finally, although these agarose wells were often sufficient to hold embryos in the right orientation for imaging, we frequently found it necessary to also use low-melting temperature agarose in conjunction with the agarose wells. Using low-melting temperature agarose can help to eliminate the occurrence of specimen drift during a long imaging session.

The accessibility of 3D-printers and ability to easily tailor the design of molds based on the specific imaging challenge without starting from scratch democratizes the ability of researchers to create customized mounting instruments for their imaging challenges. It is our hope that the molds we have designed will be useful for the zebrafish cardiac developmental biology community and the zebrafish community more generally.

## Supporting information

Supplemental data 1 - empty base designs

Supplemental data 2 - design templates

---

**Supplemental Figure 1:**
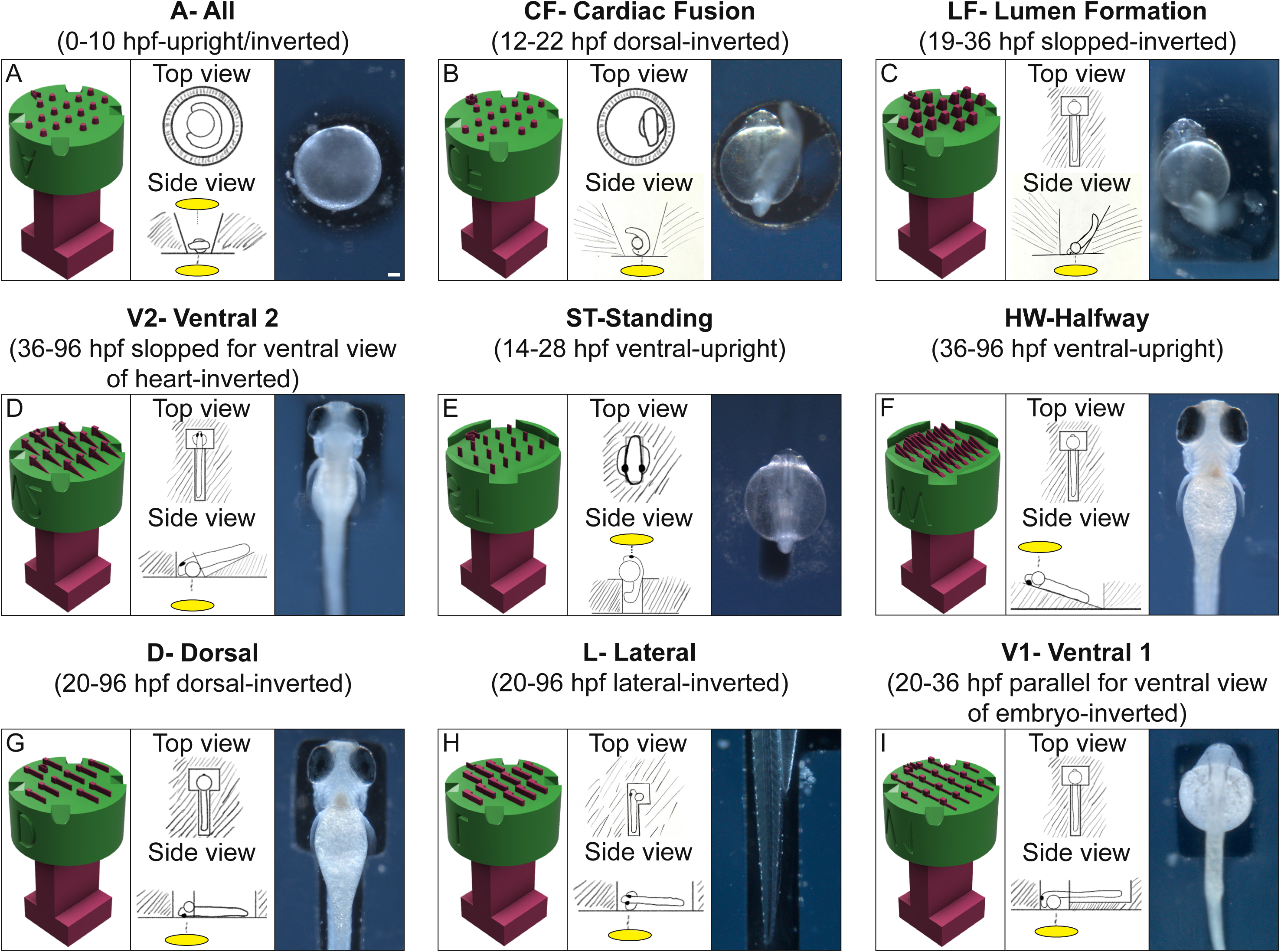
A list of molds designed to orient zebrafish embryos for imaging cardiac development and for imaging along the dorsal, lateral or ventral axes. Header for each panel contains mold name, appropriate developmental stage of embryos to mount, orientation of the embryos, and appropriate type of microscope (upright or inverted) to use for imaging. Left section of panels – design schematic, Middle section of panels – cartoon showing orientation of zebrafish embryo when mounted in an agarose well created by the mold. Yellow disc indicates location of the microscope objective. Right section of panels – picture of a zebrafish embryo mounted in an agarose well. A) Mold #A – All: this mold is designed for imaging early stages of zebrafish development (0-10 hpf) including cardiac specification using both inverted and upright microscopes. B) Mold #CF – Cardiac fusion: this mold is designed for imaging cardiac fusion (12-22 hpf) with an inverted microscope. C) Mold #LF – Lumen Formation: this mold is designed for imaging lumen formation and heart tube elongation (19-36 hpf) with an inverted microscope. D) Mold #V2 – Ventral 2: this mold is designed for imaging later cardiac developmental events such as chamber formation and cardiac maturation (36-96 hpf) with an inverted microscope. E) Mold #ST – Standing: this mold is designed for imaging the early stages of cardiac development (14-28 hpf) using an upright microscope. F) Mold #HW – Halfway: this mold is designed for imaging later cardiac developmental events with an upright microscope. G) Mold #D – Dorsal: this mold is designed for imaging the dorsal side of zebrafish embryos, including the dorsal regions of the brain. H) Mold #L – Lateral: this mold is designed for imaging the lateral regions of the zebrafish embryo. I) Mold #V1 – Ventral 1: this mold is designed for imaging the ventral side of zebrafish embryos.

**Supplemental Figure 2:**
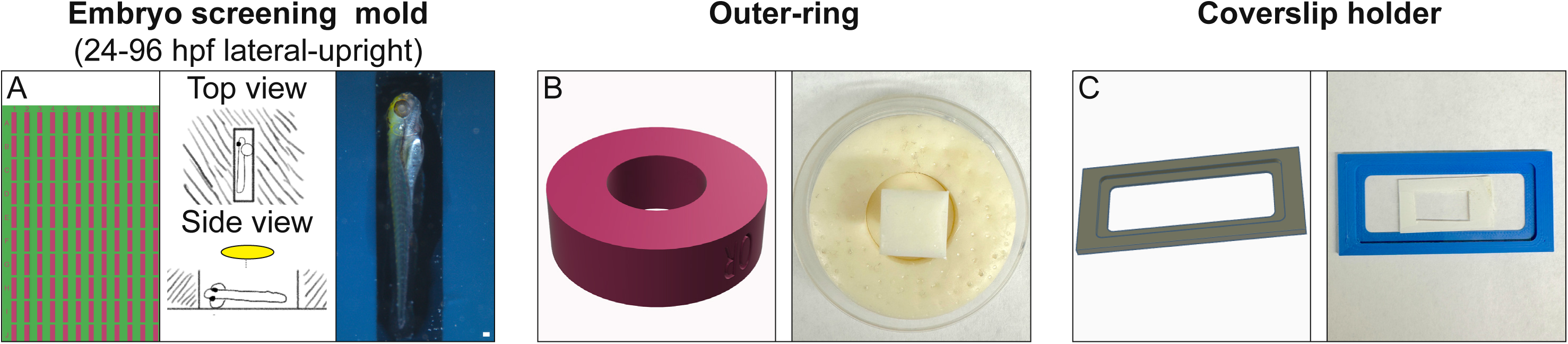
3D-printed accessories for the mounting of zebrafish embryos. **A)** A screening mold for high-throughput screening of zebrafish embryos using an upright microscope. Left section of panel - design schematic, middle section of panel - cartoon showing orientation of zebrafish embryo in an individual well, right section of panel - picture of embryo in an individual well. Scale = 100 μm **B)** The outer ring is designed to be used in conjunction with the 3D-printed inner molds (see Fig. 2) to facilitate the removal of the inner mold from the agarose wells. Left section of panel – design schematic, right section of panel – picture of outer ring and a mold prior to removal of the mold from the agarose wells. **C)** Coverslip holder to facilitate the imaging of embryos mounted using the traditional technique of placing an embryo between two coverslips. Left section of panel – design schematic, right section of panel – picture of slide holder in which layers of vinyl tape placed on a large coverslip provide a window for mounting an embryo and for preventing crushing when placing a smaller coverslip on top.

**Supplemental Figure 3:**
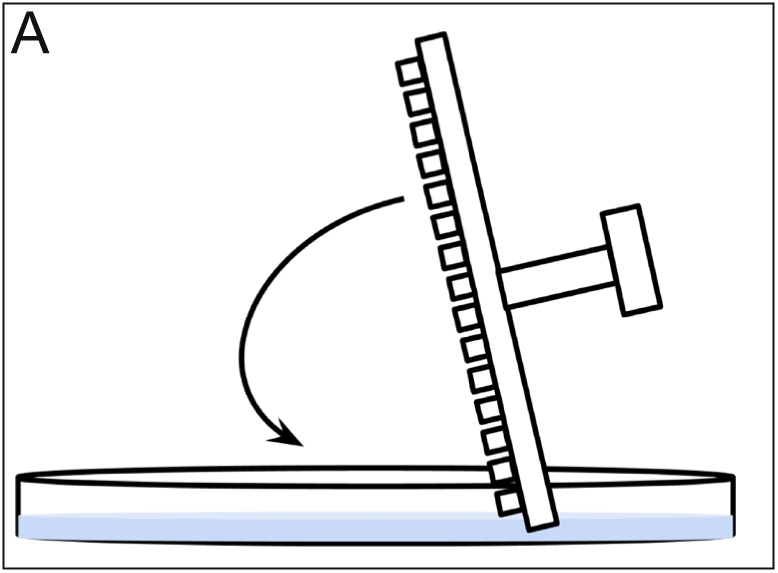
A tilting motion is used for creating agarose wells with the screening mold. **A)** Cartoon of the tilting motion needed to create agarose wells with the screening mold. Placing one side of the screening mold down and then gradually placing the rest of the mold into the petri dish prevents the introduction of bubbles.

**Supplemental movie 1: A representative time-lapse video of intersomitic vessel (ISV) formation.** Intersomitic vessel formation (ISV) was captured by using the L mold to create agarose wells in which *Tg(fli1a:egfp)* zebrafish embryos are mounted in a lateral position. Time-lapse videos were created from three-dimenstional reconstructions of confocal slices taken at 2:56 min intervals for ∼3.5 hrs, beginning at 24 hpf. Scale = 10 μm. Arrows indicate position of endothelial protrusions.

**Supplemental files A: Mold bases for inverted and upright microscopes to which pegs can be added and different pegs designs which can be further customized.**

**Supplemental files B: STL files of pre-designed molds which can be printed or modified.**

## KEY RESOURCE TABLE

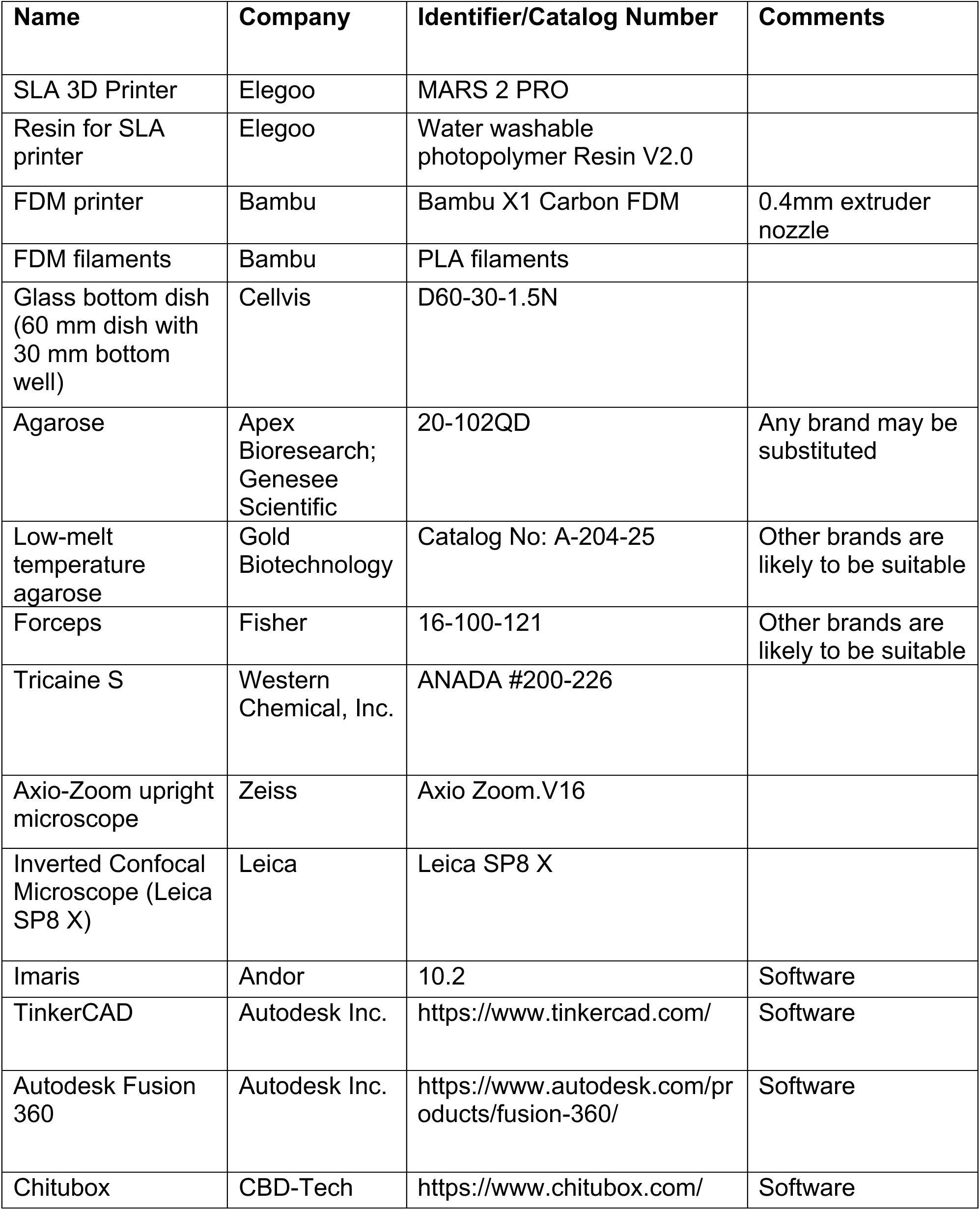
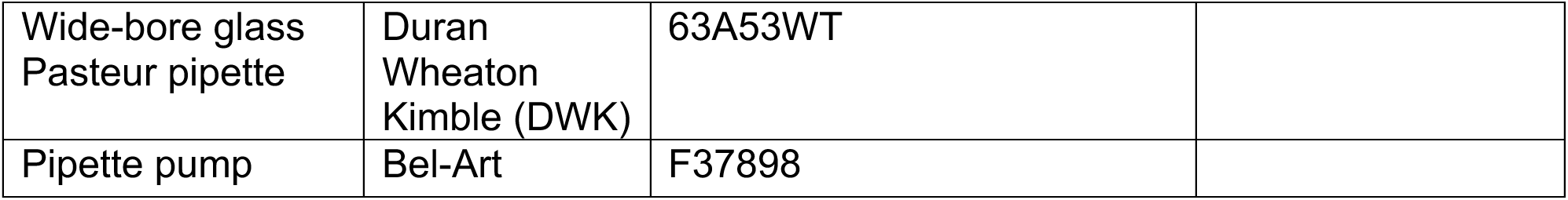

## ACKNOWLEDGMENTS

We would like to thank members of the Bloomekatz laboratory, the GlyCORE imaging core (supported by P20GM130460), the University of Mississippi IDEA lab, Dr. Yiwei Han and the Han laboratory, and members of the University of Mississippi Glycoscience and Developmental Biology communities. This work is supported by funding from the NIH (R15HD108782, P20GM130460).

